# Fat Mass Regain in Middle-Aged Mice after Sleeve Gastrectomy is divergent from plasma leptin levels

**DOI:** 10.1101/2020.10.05.326785

**Authors:** Ana BF Emiliano, Ying He, Sei Higuchi, Rabih Nemr, Natalie Lopatinsky, Gary J. Schwartz

## Abstract

**Background:** Some degree of weight regain is typically observed in human patients who undergo Sleeve Gastrectomy (SG), even if the majority of them do not return to their presurgical body weight. Although the majority of bariatric surgery patients are middle aged, most preclinical models of bariatric surgery utilize juvenile male mice. A long-term characterization of the response of mature, wild type, obese male mice to SG has not been performed.

**Methods:** Eight-month old C57bl/6J obese male mice were randomized to undergo SG, sham surgery without caloric restriction (SH) or sham surgery with caloric restriction to match body weight to the SG group (SWM). Body weight, body composition and glucose tolerance were matched at baseline. Mice were followed for 60 days following their respective surgeries.

**Results:** SG mice had a more pronounced percent weight loss than the SH group in the first post-operative month (p<0.05), along with fat mass loss (p<0.01). By the second post-operative month, the SG group started to regain fat mass, although it continued to be statistically lower than the SH group (p<0.05). Cumulative food intake was significantly lower in the SG group compared to SH group only in the first post-operative week (p<0.05), with both groups having similar cumulative food intake thereafter (p>0.05). SWM group had a significantly lower cumulative food intake throughout the study, except for week 1 (p<0.01). Glucose tolerance was only demonstrably better in the SG group compared to SH group at 8 weeks post-operatively (p<0.01). Plasma leptin was significantly lower in the SG group compared to both SWM and SH groups group by the second post-operative month (p<0.01), in spite of SG’s increasing fat mass accumulation. In the second post-operative month, both FGF-21 and GDF-15 were increased in the SH group compared to the SG and SWM groups (p<0.05), while there was no difference in plasma insulin among the three groups. Heat production was surprisingly higher in the SH group compared to the other two groups (p<0.05), even though brown adipose tissue Peroxisome Proliferator-Activated Receptor Gamma (PPARg) and Cidea mRNA expression were significantly higher in SG and SWM compared to SH (p<0.01). There was no change in BAT UCP-1 mRNA expression among the groups (p>0.05). There was also no change in fecal lipid content among the groups (p>0.05).

**Conclusions:** SG in obese, middle aged male mice leads is accompanied by fat mass regain in the second post-operative month, while plasma leptin levels continue to be significantly lower. This raises the question of whether the observed fat mass regain consists mostly of visceral adipose tissue.

## Introduction

Weight regain is a common phenomenon that occurs between 2-7 years after Sleeve Gastrectomy (SG), with at least 20 percent of patients regaining enough weight to require a new bariatric operation^1,2,3,4^. The exact percentage of patients that regain weight after SG is not known, as a result of lack of standardization of weight regain and poor reporting^5^. It is now known what leads to weight regain after SG. One study of human subjects indicated that both fat and lean mass are lost initially after bariatric surgery, while one year post-operatively, most of the loss represent fat mass^6^. The implications of weight regain are numerous, including the need for reoperation in order to prevent obesity and type 2 diabetes recidivism^7,8^.

Rodent models of bariatric surgery are invaluable tools in the discovery of mechanistic underpinnings of the beneficial metabolic effects of bariatric surgery^9,10,11^. However, one of the weaknesses of these models is the almost exclusive utilization of juvenile male mice. The mean age of bariatric surgery patients is approximately 45-50^12,13,14^. Although insight acquired from preclinical rodent models can be extremely useful, there are limitations when these models do not reflect the human patient population that the study is targeting, whether adolescents, young adults, middle aged or older adults. Age is an important factor in the clinical response to therapeutic interventions in humans. It would be surprising if mouse age did not influence outcomes in rodent bariatric surgery studies. Moreover, the chronicity of exposure to a high fat diet may also be significant in the mouse response to bariatric surgery. Therapeutic responses from mice fed a high fat diet for 8 weeks may not be comparable to the response from mice fed a high fat diet for 16 weeks or longer. Intuitively, one would estimate that mice with a shorter time exposure to high fat diet would have a better metabolic response to SG, with increased weight loss and lower blood glucose.

The present study’s goal was to evaluate weight regain in a middle aged, obese male mouse model. DIO (diet-induced obese) mice on a 60% high fat diet, started at 6 weeks of age were used for the experiments described here. At 8 months old, these mice were randomized into SG group, sham surgery with ad libitum diet (SH) or sham surgery weight-matched (SWM) to the SG group, through caloric restriction. We found that these older, mature obese mice responded did not have a homogenous response to SG, with some of them losing much greater percentages of weight than others, in addition to maintaining that weight loss. Although by the end of the study they had a similar body weight as the SWM group, their caloric intake was higher. Glucose tolerance only improved in the intraperitoneal glucose tolerance test, with no improvement in oral glucose tolerance or in insulin tolerance. Moreover, the most intriguing finding was that of a significantly lower plasma leptin level in the SG group, in spite of a higher body weight then SWM group and in the face of increasing accumulation of fat mass in the SG group.

## Methods

### Animals

All experiments were approved and performed in accordance to the Columbia University Institutional Animal Care and Use Committee. Male C57bl/6J on a 60% high fat diet were purchased from The Jackson Laboratories at 16 weeks of age and maintained on the same high fat diet (Research Diets, catalogue number D12492, New Jersey, NJ). At 8 months of age, mice were randomized to SG, SH and SWM to the SG group (n=14 for SG; n=6 for SH, n=7 for SWM) and single-housed. This study was replicated two other times. Mice were sacked after 60 days post-operatively, after a 5 hour fast, under isoflurane anesthesia, when intracardiac blood collection and tissue harvest were performed. Body weight and food intake were measured daily for the first week post-operatively and weekly thereafter. Body composition with EchoMRI was done at baseline, at one month and two months post-operatively. Glucose tolerance tests were done at baseline, and then at 2 and 8 weeks post-operatively; insulin tolerance test was performed 4 weeks post-operatively. Feces were collected for fecal lipid content two months post-operatively. Plasma and tissue samples for mRNA analysis were also obtained 60 days post-operatively. Mice spent one week in metabolic cages between 7-8 weeks post-operatively. Baseline measurements are defined as having being taken 2 weeks prior to surgeries.

### Sleeve gastrectomy technique

All surgical procedures were sterile. Mice received meloxicam, saline and enrofloxacin subcutaneously at the time of the surgery (Boehringer Ingelheim; saline from Medline; Bayer). A midline laparotomy was performed and the stomach isolated from surrounding connective tissue. Surgeries were performed under a dissecting microscope (Leica M-125). For SG, the gastric arteries were ligated with 8-0 suture (Vicryl Violet J401G, BV130-5 Taper, Ethicon). After ligation, an incision was made at the bottom of the fundus and the stomach contents emptied. The stomach was irrigated with warm saline. A 6-0 suture (Vicryl J212H, Ethicon) was used for continuous suturing from about 2 millimeters below the esophagogastric junction, to the point where the pancreas is attached to the stomach. Tissue below that line was dissected with microscissors. The edges of the stomach where the resection occurred were sutured together with 7-0 suture (Vicryl J488G, P-1 cutting, Ethicon). Gastric leaks were assessed. Sham surgeries were as above, except that after stomach isolation, the stomach was put back into the abdominal cavity and the abdominal wall closed in layers with 5-0 suture (Vicryl J493G, Ethicon). Mice were kept on a liquid diet with Ensure High Protein (Abbott Laboratories) and high fat diet for 4 days prior to the surgeries. The day before the surgery, high fat diet was removed and the mice were kept on Ensure until the 7^th^ post-operative day. High fat diet was reintroduced on the 5^th^ post-operative day and continued until the end of the study. SG and Sham groups were kept on ad libitum high fat diet. The sham weight-matched group was kept on 2-2.5 grams of high fat diet daily, provided at different times of the day to prevent entrainment. The SG surgery described here is a modification of an SG surgical technique previously developed by another group^16^.

### Glucose Tolerance Test and Insulin Tolerance Test

Oral glucose tolerance test (OGTT) was performed by oral gavage using a plastic needle (1FTP-20-38, Instech Laboratories, Inc.), with 2 grams of dextrose per kg of body weight. Intraperitoneal glucose tolerance test was performed by injecting the mice intraperitoneally with 2 grams of dextrose per kg of body weight. Insulin Tolerance Test was performed using regular insulin at a dose of 0.75 units per Kg of body weight (Humulin R, Lilly).

### Energy expenditure

Mice were in Comprehensive Lab Animal Monitoring System (CLAMS, Columbus Instruments, Columbus, OH) for one week, between 7-8 weeks post-operatively.

### Body Composition

Body composition was measured at baseline, at one and two months post-operatively using Echo-MRI™-100H (EchoMRI LLC, Houston, TX).

### Fecal Lipid Content Assay

Feces were collected from individual mice. Feces were dried overnight on a six-well plate, at 42°C. For each reaction, 100 mg of feces (per mouse) was placed in 1 mL of NaCl and then homogenized with beads for five minutes. The solution was transferred to a 15 mL tube, to which a mix of chloroform and methanol at 2:1 was added (Chloroform, Fisher, C298-4; Methanol, Fisher A456-212). The mixture was vortexed vigorously and then centrifuged at 2000g for 10 minutes. The chloroform phase was collected and transferred to a 20 mL vial and evaporated under N2. After that, 1 mL of deionized water was added. The next day, samples were assayed in duplicates, with a Free Fatty Acid (FFA) kit (Wako 995-34693; 997-34893; Fujifilm USA). Samples were diluted at 1:20 and incubated at 37°C for five minutes and read with a microplate reader.

### RNA extraction and analysis

RNA extraction was performed using NucleoSpin RNA Set for Nucleozol (Macherey-Nagel, catalogue #740406.50). cDNA was obtained using the High Capacity cDNA Reverse Transcription Kit (Applied BioSystems, ThermoFisher Scientific, catalogue #4368813). For RT-PCR, we used ready-made probes purchased from Integrated DNA Technologies (IDT, USA). The enzyme used was Taqman Fast Advanced Master Mix (ThermoFisher Scientific, catalogue #4444557). Samples were run in triplicates and the experiment replicated three times. The instrument was QuantiStudio™ 5 Real Time PCR system (ThermoFisher Scientific). We used the QuantiStudio™ 5 software to perform the analysis.

### Plasma Assays

Blood was collected from animals fasted for 5 hours, via intracardiac puncture. Blood was collected in tubes containing EDTA, aprotinin and Diprotin A, a DPPIV inhibitor (EDTA, ThermoFisher Scientific, catalogue #15575020; Aprotinin, Millipore Sigma, catalogue #A1153; Diprotin A, Enzo Scientific, catalogue #ALX-260-036-M005). Blood was spun for 15 min at 2000 g, in a refrigerated benchtop microcentrifuge and subsequently stored in a minus 80 freezer. For FGF-21, Leptin and Insulin plasma measurement, we used a U-PLEX MSD assay (MesoScaleDiscovery, catalogue #K152ACL-1). GDF-15 was measured using an ELISA kit (R&D Biosystems, catalogue #MGD150).

### Statistical Analysis

We used GraphPad Prism 8.0 version software for all statistical analysis. One Way ANOVA was used for comparison of three groups. Two-Way ANOVA with repeated measures was used for comparisons of three groups with multiple measures. XY analysis was used for generating AUC graphs used for glucose tolerance and energy expenditure analysis, which were subsequently analyzed with ANOVA. Power was set at 80%, with a p value of <0.05.

## Results

### Body Weight

SG and SWM had body weight, fat mass and lean mass that were not statistically different at baseline (p>0.05). SH had a slightly lower body weight at baseline compared to SG and SWM (p<0.05), but similar fat and lean mass compared to the SG and SWM groups at baseline (p>0.05) (figure 1, top panel). Average body weight at baseline was 45 grams. One month after surgeries, both SG and SWM had statistically lower body weight (p<0.05) and fat mass (p<0.01), compared to the sham group (figure 1, middle panel). Lean mass was not different among groups (p>0.05). At two months, SG and SWM continued to have lower body weight (p<0.05) and fat mass than SH (p<0.05), but to a smaller degree (figure 1, bottom panel). Lean mass was not different among groups at 2 months, either (p>0.05). Percent weigh loss significantly lower in SG and SWM both in the first month and second months post-operatively (p<0.01) (figure 2). However, in the second month, half of the SG subjects had regained at least half of their lost weight back. The overall statistics do not reflect this finding because a few subjects in the SG group had very pronounced weight loss. Fecal lipid content was not different among groups (p>0.05) (figure 2).

**Figure 1.**
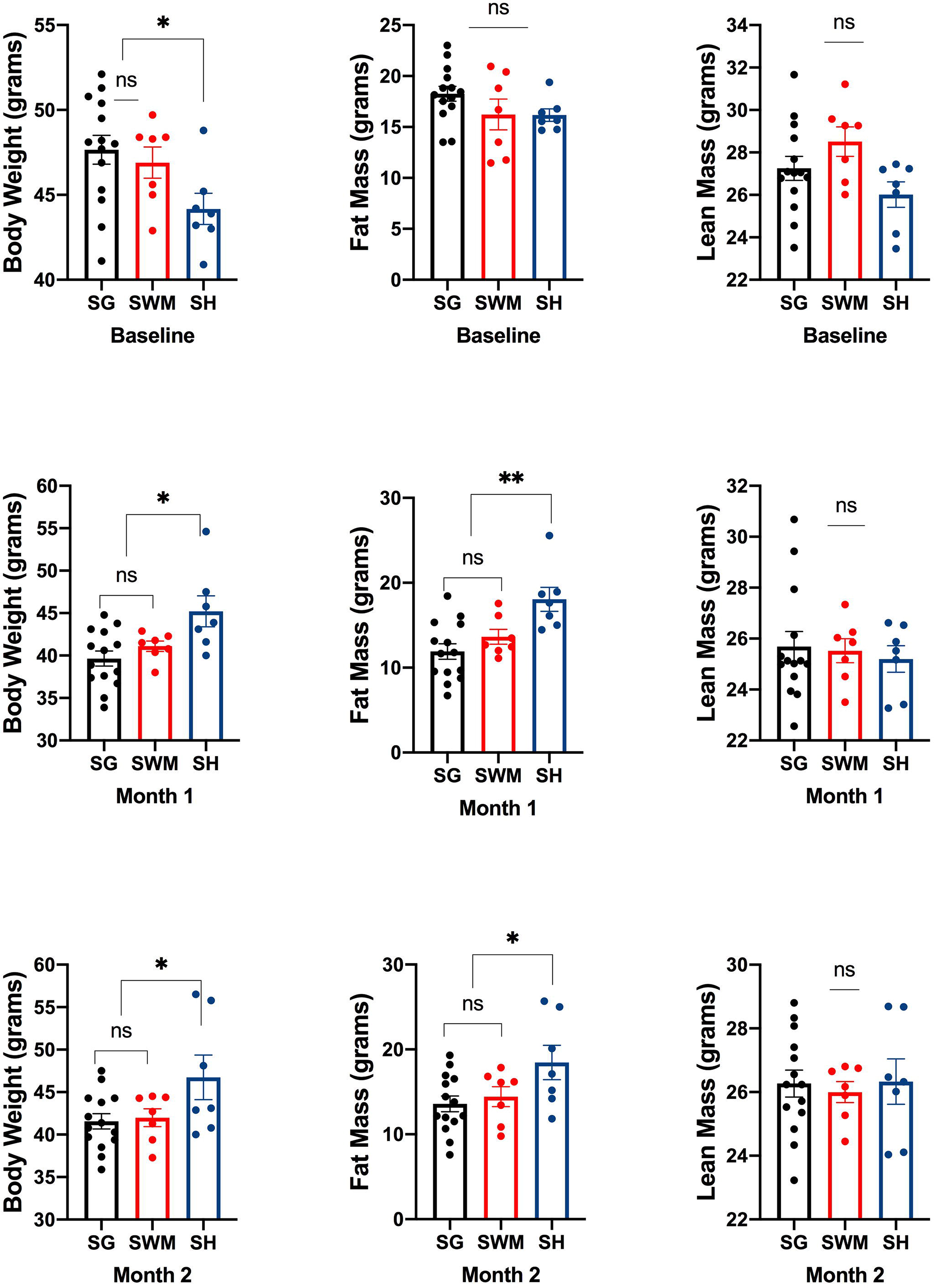
Top panel depicts body weight and body composition at baseline. Middle panel depicts body weight and body composition in the first month after surgeries. Bottom panel shows body weight and body composition in the second month after surgeries. SG – sleeve gastrectomy group; SWM – sham weight matched; SH – sham. NS – nonsignificant; * p<0.05; ** p<0.01. For n and statistical analysis, please see text.

**Figure 2.**
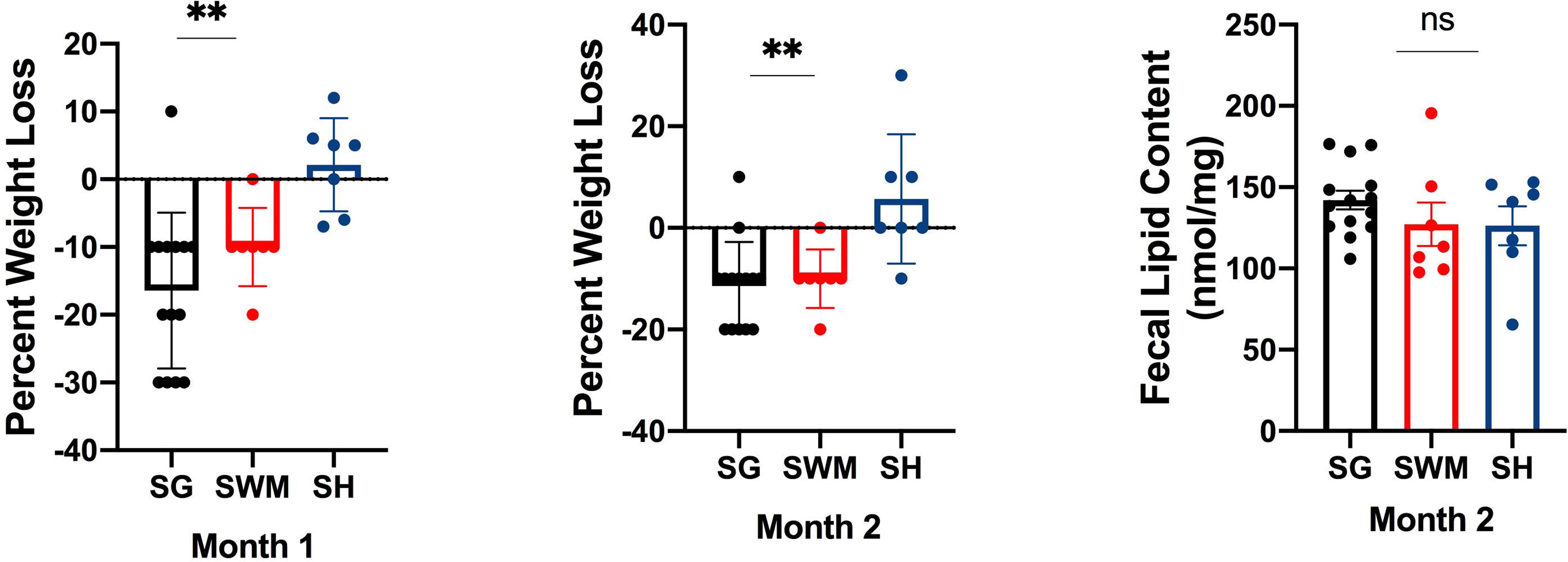
First two panels show percent weight loss at 1 and 2 months after surgeries. Last panel shows fecal lipid content. SG – sleeve gastrectomy group; SWM – sham weight matched; SH – sham. NS – nonsignificant; * p<0.05; ** p<0.01. For n and statistical analysis, please see text.

### Food Intake

As previously reported, SG mice had a significantly lower cumulative food intake in the first post-operative week, compared to the SH group (p<0.05) (figure 3)^17^. SG had a similar cumulative food intake as the SH group throughout the rest of the study, after the first post-operative week (p>0.05) (figure 3). SWM had a significantly lower cumulative food intake than SG and SH groups in the first and second post-operative months (p<0.01) (figure 3).

**Figure 3.**
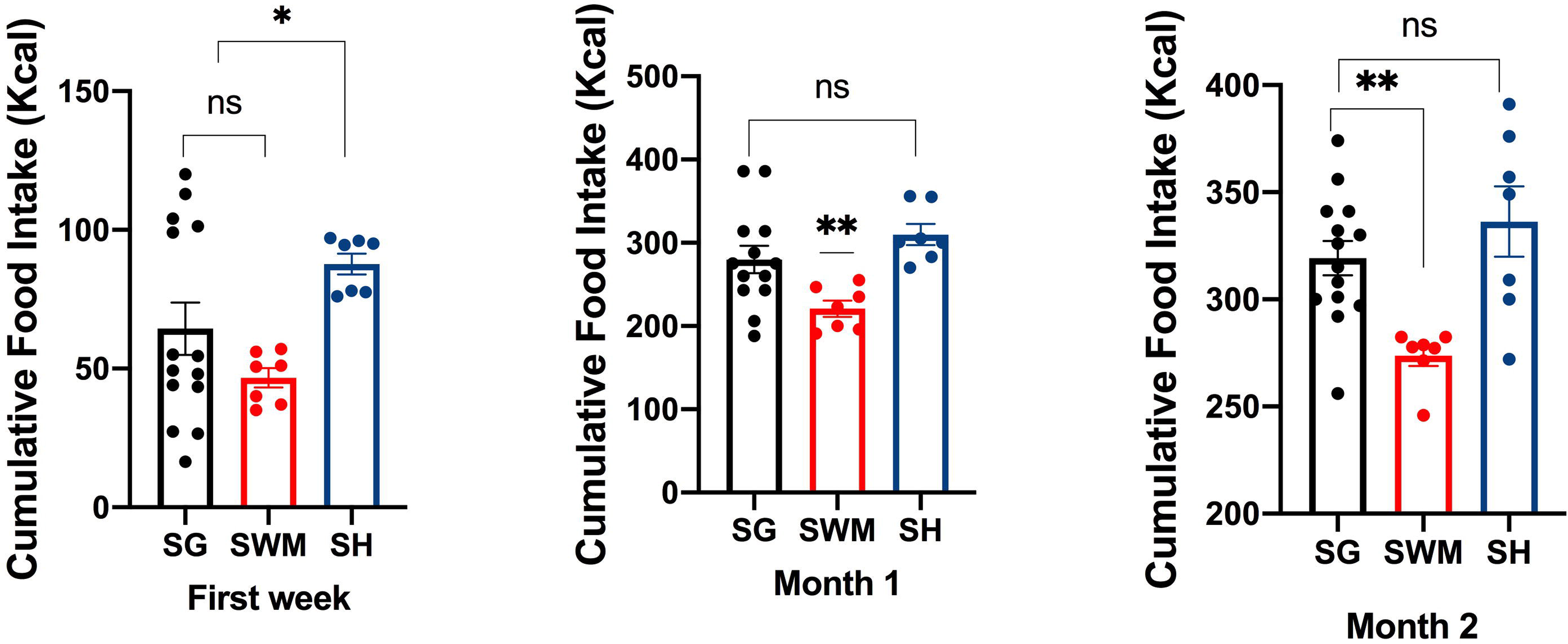
Cumulative food intake at the first post-operative week and then at 1 and 2 months after surgeries. SG – sleeve gastrectomy group; SWM – sham weight matched; SH – sham. NS – nonsignificant; * p<0.05; ** p<0.01. For n and statistical analysis, please see text.

### Glucose Tolerance and Insulin Tolerance

IPGTT at baseline did not reveal any differences in AUC among the three groups, with all groups being hyperglycemic (p>0.05) (figure 4, top panel). OGTT in the second week post-operatively, did not show a significantly lower AUC for SG compared to both SWM and SH (p>0.05) (figure 4, second panel). An ITT at the fourth post-operative week revealed no differences in AUC among the groups (p<0.05) (figure 4, third panel). At 8 weeks post-operatively, the IPGTT AUC was significantly lower in the SG and SWM groups compared to the SWM and SH groups (p<0.05) (figure 5, top panel). At 8 weeks post-operatively, the OGTT AUC was significantly lower in the SG and SWM groups, compared to SH control (p<0.01) (figure 4, bottom panel).

**Figure 4.**
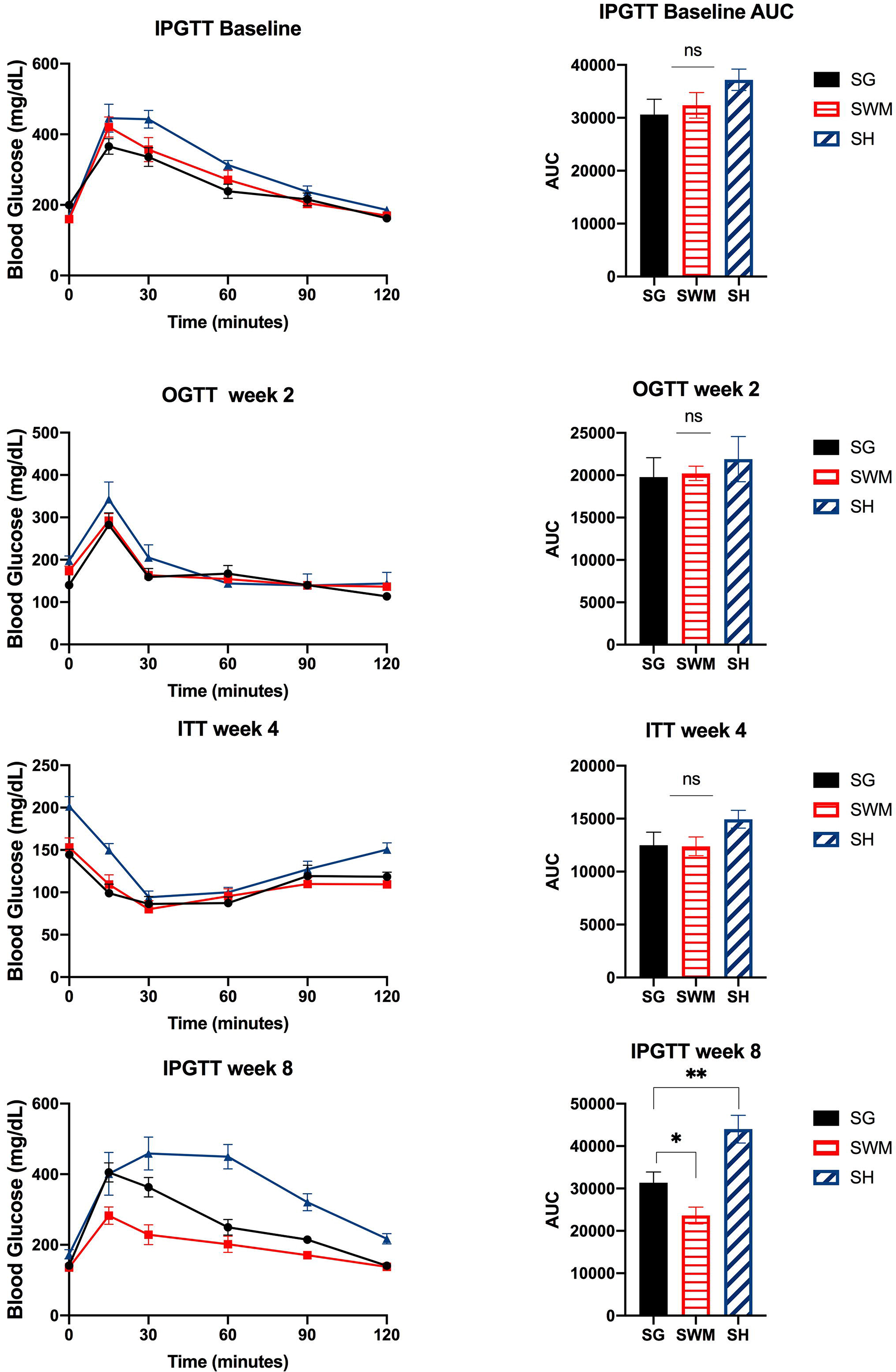
Top panel shows baseline IPGTT and IPGTT AUC. Second panel shows OGTT and OGTT AUC 2 weeks post-operatively. Third panel shows ITT and ITT AUC 4 weeks post-operatively. Bottom panel shows IPGTT and IPGTT AUC 8 weeks post-operatively. SG – sleeve gastrectomy group; SWM – sham weight matched; SH – sham. NS – nonsignificant; * p<0.05; ** p<0.01. For n and statistical analysis, please see text. IPGTT-intraperitoneal glucose tolerance test; OGTT – oral glucose tolerance test; ITT – Insulin Tolerance Test.

**Figure 5.**
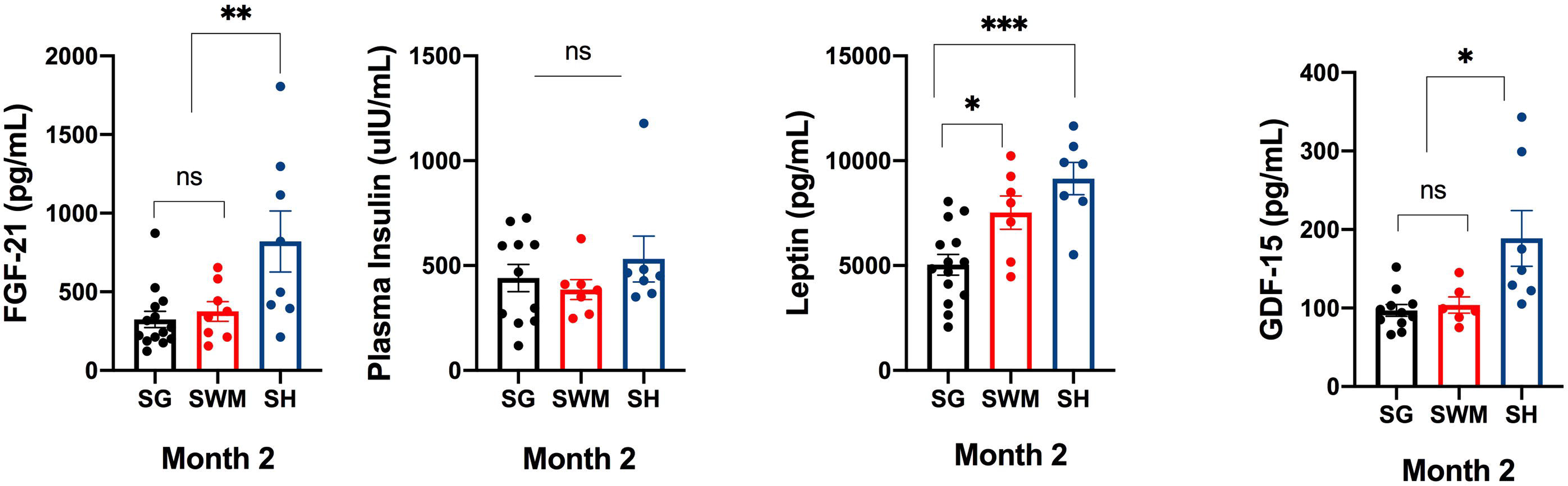
Plasma levels of FGF-21, insulin, leptin and GDF-15. SG – sleeve gastrectomy group; SWM – sham weight matched; SH – sham. NS – nonsignificant; * p<0.05; ** p<0.01. For n and statistical analysis, please see text.

### Plasma hormones

At the end of the second month post-operatively, plasma FGF-21 and GDF-15 were significantly higher in the SH group, compared to the SG and SWM groups (p<0.01 for FGF-21 and p<0.05 for GDF-15) (figure 5). Plasma insulin was not different among the groups (p>0.05). Plasma leptin was significantly lower in the SG group compared to the SWM and SH groups (p<0.05 for comparison with SWM group and p<0.001 for comparison with SH group).

### Energy expenditure and BAT mRNA expression

The SG and the SWM groups had an increased mRNA expression of genes that regulate metabolic activity in BAT, namely Peroxisome proliferator-activated receptor gamma (PPARgamma) and Cidea (p<0.01) (figure 6, top panel). Uncoupling Protein 1(UCP-1) mRNA expression was not different among the three groups (p>0.05) (figure 8, top panel). Surprisingly, heat production was slightly higher in the SH group compared to the SG and SWM groups (p<0.05) (figure 6, middle panel). Respiratory Exchange Ratio (RER) was not different among the three groups (p>0.05) (figure 6, bottom panel).

**Figure 6.**
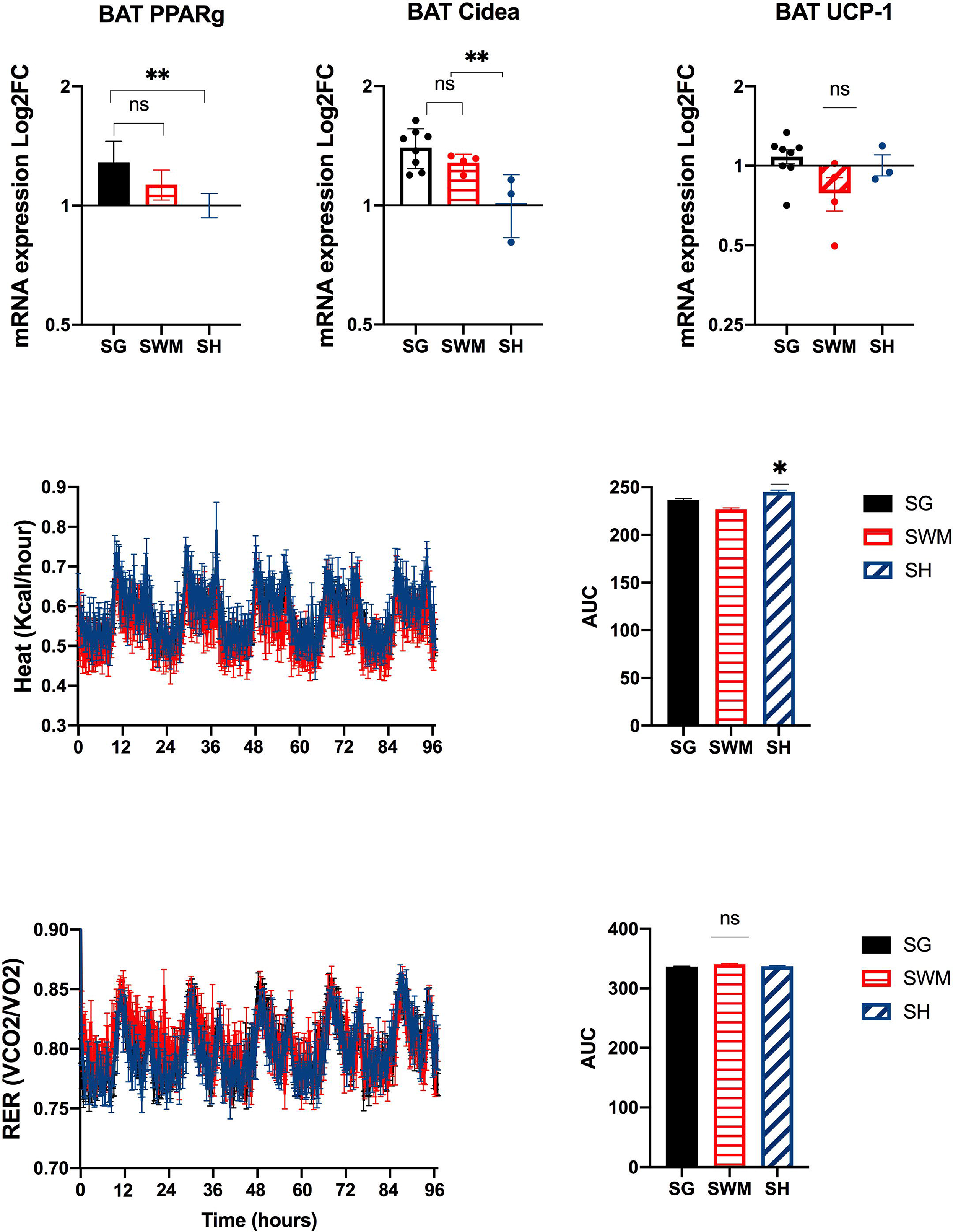
Top panel shows mRNA expression in BAT, including PPAR gamma, Cidea and UCP-1. Middle panel shows heat production over time. Bottom panel shows RER over time. SG – sleeve gastrectomy group; SWM – sham weight matched; SH – sham. Peroxisome proliferator-activated receptor gamma – PPARgamma; Uncoupling Protein 1 – UCP-1. NS – nonsignificant; * p<0.05; ** p<0.01; ***p<0.001; ****p<0.0001. For n and statistical analysis, please see text.

## Discussion

Approximately one third of American adults are obese^17^. For individuals with a BMI higher than 35, the lifetime risk of developing type 2 diabetes is estimated at 70%^18^. Severe obesity, defined as a BMI greater than 40 has been projected to double within the next two decades^19^. Consequently, it is likely that the demand for bariatric surgeries such as SG will only increase. Most individuals with obesity and type 2 diabetes lack access to bariatric surgery, with only 2% of all eligible patients in the US undergoing these procedures^20^. A more complete understanding of the mechanisms whereby SG promotes durable weight loss and blood glucose clinical improvement is necessary for the development of more accessible, non-invasive therapies for obesity and type 2 diabetes. Likewise, a better understanding of weight regain or failure to lose weight after SG can lead to important insights into mammalian energy homeostasis. In addition, it may also pave the way for the development of more durable and effective therapies for type 2 diabetes and obesity.

In the present study, we found that middle-aged, obese male mice do not respond to SG uniformly, with 50% of mice regaining weight, and with 10% of mice in the SG group never losing any weight, in spite of the surgeries having been performed by the same surgeon, using the exact same technique. Mice in the SG group started to gain fat mass after the first post-operative month, until the time when the study was ended. A longer study may have uncovered a trajectory of fat mass regain, which could have ultimately lead to no differences in body weight between the SG and the SH group. It was intriguing that plasma leptin was significantly lower in the SG group, even when compared to the SWM group that had the same body weight and fat mass as the SG group. It has been reported that plasma leptin levels correlate more closely with subcutaneous fat depots than visceral fat^21,22^. It is plausible that the significantly lower plasma leptin in the SG group reflects an increase in visceral fat as opposed to accumulation of subcutaneous fat. Future studies should use imaging methodologies such as MRI to explore differences in the distribution of adipose tissue compartments after SG, especially in weight regain.

The trend toward weight regain observed in the SG group may be linked to failure of the surgery to suppress food intake^16^. Long-term studies of bariatric surgery outcomes have found that a sustained suppression of food intake is associated with better outcomes in terms of weight loss, glucose homeostasis and cardiovascular risk reduction^23,24^. Future studies should also focus on understanding the nature of the self-limited anorexia phase after bariatric surgery in rodent models, as this would advance our knowledge of gut-brain interactions. Also, designing studies that would permit analysis of subjects in mouse cohorts that continue to maintain a lower food intake. That type of analysis was not possible in this study because only one mouse in the SG cohort continued to have a lower food intake. This observation was made in the three mouse cohorts that we followed for this manuscript.

We did not observe a significant improvement in oral glucose tolerance or insulin tolerance. Intraperitoneal glucose tolerance was only statistically lower in the SG group compared to the SH group at 8 weeks post-operatively. It is not clear why. Compared to the baseline IPGTT, all groups displayed improved glucose homeostasis during the OGTT at week 2, although these two tests cannot be directly compared. At week 8, it looks like the SG had the same performance as at baseline, with SWM being the only group that truly manifested an improved glucose homeostasis. This underscores the importance of dietary intake in blood glucose regulation, since the SWM was the only group on caloric restriction, having a significantly lower cumulative food intake compared to the other two groups.

In spite of the literature associating FGF-21 to increased peripheral glucose uptake and improved glucose homeostasis, it was the SH group that had increased FGF-21 plasma levels^25^. An increase in fasting and post-prandial FGF-21 was observed in adolescents up to three months post SG^26^. Additionally, FGF-21 was found to correlate with weight loss in adolescents following SG^27^. We wonder if the lack of an increase in FGF-21 in SG in our study was due to the age of the mice, or whether we missed an early increase in FGF-21 by only testing plasma samples from 2 months after SG. Deletion of Growth Differentiation Factor-15 (GDF-15) was not found to affect food intake or body weight after SG^28^. But in humans, plasma levels of GDF-15 were found to positively correlated with weight loss magnitude^29^.

In terms of energy expenditure, we did not find evidence of increased energy expenditure in SG mice. It is not clear what prevents more pronounced weight gain in the SG group, considering that their cumulative food intake is similar as to the SH group. It is unlikely that the significant increase in the BAT mRNA expression of PPARgamma and Cidea are responsible for preventing more significant weight gain in the SG group.

In summary, we found that middle-aged, obese and glucose intolerant male mice have a heterogeneous response to SG in terms of weight loss, with 50% quickly regaining fat mass at 2 months post SG. Their glycemic response to SG was also limited, which underscores the important of dietary intake for glucose homeostasis. We believe their older age and a prolonged, chronic exposure to high fat diet prevented these mice from being more responsive to the beneficial metabolic effects of SG.

## References

1. Lauti M, Kularatna M, Hill AG, MacCormick AD. Weight Regain Following Sleeve Gastrectomy-a Systematic Review. Obes Surg. 2016 Jun;26(6):1326–34. doi: 10.1007/s11695-016-2152-x. PMID: 27048439.

2. Himpens J, Dobbeleir J, Peeters G. Long-term results of laparoscopic sleeve gastrectomy for obesity. Ann Surg. 2010 Aug;252(2):319–24. doi: 10.1097/SLA.0b013e3181e90b31. PMID: 20622654.

3. Kushner RF, Sorensen KW. Prevention of Weight Regain Following Bariatric Surgery. Curr Obes Rep. 2015 Jun;4(2):198–206. doi: 10.1007/s13679-015-0146-y. PMID: 26627215.

4. Fahmy MH, Sarhan MD, Osman AM, Badran A, Ayad A, Serour DK, Balamoun HA, Salim ME. Early Weight Recidivism Following Laparoscopic Sleeve Gastrectomy: A Prospective Observational Study. Obes Surg. 2016 Nov;26(11):2654–2660. doi: 10.1007/s11695-016-2165-5. PMID: 27056195.

5. Lauti M, Lemanu D, Zeng ISL, Su’a B, Hill AG, MacCormick AD. Definition determines weight regain outcomes after sleeve gastrectomy. Surg Obes Relat Dis. 2017 Jul;13(7):1123–1129. doi: 10.1016/j.soard.2017.02.029. Epub 2017 Mar 9. PMID: 28438493.

6. Maïmoun L, Lefebvre P, Aouinti S, Picot MC, Mariano-Goulart D, Nocca D; Montpellier Study Group of Bariatric Surgery. Acute and longer-term body composition changes after bariatric surgery. Surg Obes Relat Dis. 2019 Nov;15(11):1965–1973. doi: 10.1016/j.soard.2019.07.006. Epub 2019 Jul 29. PMID: 31519485.

7. Switzer NJ, Karmali S, Gill RS, Sherman V. Revisional Bariatric Surgery. Surg Clin North Am. 2016 Aug;96(4):827–42. doi: 10.1016/j.suc.2016.03.004. PMID: 27473804.

8. Shah A, Laferrère B. Diabetes after Bariatric Surgery. Can J Diabetes. 2017 Aug;41(4):401–406. doi: 10.1016/j.jcjd.2016.12.009. Epub 2017 Apr 27. PMID: 28457649; PMCID: PMC5875725.

9. Ryan KK, Tremaroli V, Clemmensen C, Kovatcheva-Datchary P, Myronovych A, Karns R, Wilson-Pérez HE, Sandoval DA, Kohli R, Bäckhed F, Seeley RJ. FXR is a molecular target for the effects of vertical sleeve gastrectomy. Nature. 2014 May 8;509(7499): 183–8. doi: 10.1038/nature13135. Epub 2014 Mar 26. PMID: 24670636; PMCID: PMC4016120.

10. Saeidi N, Meoli L, Nestoridi E, Gupta NK, Kvas S, Kucharczyk J, Bonab AA, Fischman AJ, Yarmush ML, Stylopoulos N. Reprogramming of intestinal glucose metabolism and glycemic control in rats after gastric bypass. Science. 2013 Jul 26;341(6144):406–10. doi: 10.1126/science.1235103. PMID: 23888041; PMCID: PMC4068965.

11. McGavigan AK, Garibay D, Henseler ZM, Chen J, Bettaieb A, Haj FG, Ley RE, Chouinard ML, Cummings BP. TGR5 contributes to glucoregulatory improvements after vertical sleeve gastrectomy in mice. Gut. 2017 Feb;66(2):226–234. doi: 10.1136/gutjnl-2015-309871. Epub 2015 Oct 28. PMID: 26511794; PMCID: PMC5512436.

12. Arterburn D, Wellman R, Emiliano A, Smith SR, Odegaard AO, Murali S, Williams N, Coleman KJ, Courcoulas A, Coley RY, Anau J, Pardee R, Toh S, Janning C, Cook A, Sturtevant J, Horgan C, McTigue KM; PCORnet Bariatric Study Collaborative. Comparative Effectiveness and Safety of Bariatric Procedures for Weight Loss: A PCORnet Cohort Study. Ann Intern Med. 2018 Dec 4;169(11):741–750. doi: 10.7326/M17-2786. Epub 2018 Oct 30. PMID: 30383139; PMCID: PMC6652193.

13. Courcoulas AP, King WC, Belle SH, Berk P, Flum DR, Garcia L, Gourash W, Horlick M, Mitchell JE, Pomp A, Pories WJ, Purnell JQ, Singh A, Spaniolas K, Thirlby R, Wolfe BM, Yanovski SZ. Seven-Year Weight Trajectories and Health Outcomes in the Longitudinal Assessment of Bariatric Surgery (LABS) Study. JAMA Surg. 2018 May 1;153(5):427–434. doi: 10.1001/jamasurg.2017.5025. PMID: 29214306; PMCID: PMC6584318.

14. Buchwald H, Avidor Y, Braunwald E, Jensen MD, Pories W, Fahrbach K, Schoelles K. Bariatric surgery: a systematic review and meta-analysis. JAMA. 2004 Oct 13;292(14):1724–37. doi: 10.1001/jama.292.14.1724. Erratum in: JAMA. 2005 Apr 13;293(14):1728. PMID: 15479938.

15. Garibay D, Cummings BP. A Murine Model of Vertical Sleeve Gastrectomy. J Vis Exp. 2017 Dec 18;(130):56534. doi: 10.3791/56534. PMID: 29286478; PMCID: PMC5755614.

16. Mumphrey MB, Hao Z, Townsend RL, Patterson LM, Münzberg H, Morrison CD, Ye J, Berthoud HR. Eating in mice with gastric bypass surgery causes exaggerated activation of brainstem anorexia circuit. Int J Obes (Lond). 2016 Jun;40(6):921–8. doi: 10.1038/ijo.2016.38. Epub 2016 Mar 17. PMID: 26984418; PMCID: PMC4899289.

17. Ogden CL, Carroll MD, Kit BK, Flegal KM. Prevalence of childhood and adult obesity in the United States, 2011-2012. JAMA. 2014 Feb 26;311(8):806–14. doi: 10.1001/jama.2014.732. PMID: 24570244; PMCID: PMC4770258.

18. Narayan KM, Boyle JP, Thompson TJ, Gregg EW, Williamson DF. Effect of BMI on lifetime risk for diabetes in the U.S. Diabetes Care. 2007 Jun;30(6): 1562–6. doi: 10.2337/dc06-2544. Epub 2007 Mar 19. PMID: 17372155.

19. Finkelstein EA, Khavjou OA, Thompson H, Trogdon JG, Pan L, Sherry B, Dietz W. Obesity and severe obesity forecasts through 2030. Am J Prev Med. 2012 Jun;42(6):563–70. doi: 10.1016/j.amepre.2011.10.026. PMID: 22608371.

20. 6.English WJ, DeMaria EJ, Brethauer SA, Mattar SG, Rosenthal RJ, Morton JM. American Society for Metabolic and Bariatric Surgery estimation of metabolic and bariatric procedures performed in the United States in 2016. Surg Obes Relat Dis. 2018 Mar;14(3):259–263. doi: 10.1016/j.soard.2017.12.013. Epub 2017 Dec 16. PMID: 29370995.

21. Russell CD, Petersen RN, Rao SP, Ricci MR, Prasad A, Zhang Y, Brolin RE, Fried SK. Leptin expression in adipose tissue from obese humans: depot-specific regulation by insulin and dexamethasone. Am J Physiol. 1998 Sep;275(3):E507–15. doi: 10.1152/ajpendo.1998.275.3.E507. PMID: 9725819.

22. Van Harmelen V, Reynisdottir S, Eriksson P, Thörne A, Hoffstedt J, Lönnqvist F, Arner P. Leptin secretion from subcutaneous and visceral adipose tissue in women. Diabetes. 1998 Jun;47(6):913–7. doi: 10.2337/diabetes.47.6.913. PMID: 9604868.

23. Sjöström L, Lindroos AK, Peltonen M, Torgerson J, Bouchard C, Carlsson B, Dahlgren S, Larsson B, Narbro K, Sjöström CD, Sullivan M, Wedel H; Swedish Obese Subjects Study Scientific Group. Lifestyle, diabetes, and cardiovascular risk factors 10 years after bariatric surgery. N Engl J Med. 2004 Dec 23;351(26):2683–93. doi: 10.1056/NEJMoa035622. PMID: 15616203.

24. Laurenius A, Larsson I, Bueter M, Melanson KJ, Bosaeus I, Forslund HB, Lönroth H, Fändriks L, Olbers T. Changes in eating behaviour and meal pattern following Roux-en-Y gastric bypass. Int J Obes (Lond). 2012 Mar;36(3):348–55. doi: 10.1038/ijo.2011.217. Epub 2011 Nov 29. PMID: 22124454.

25. Kharitonenkov A, Shiyanova TL, Koester A, Ford AM, Micanovic R, Galbreath EJ, Sandusky GE, Hammond LJ, Moyers JS, Owens RA, Gromada J, Brozinick JT, Hawkins ED, Wroblewski VJ, Li DS, Mehrbod F, Jaskunas SR, Shanafelt AB. FGF-21 as a novel metabolic regulator. J Clin Invest. 2005 Jun;115(6):1627–35. doi: 10.1172/JCI23606. Epub 2005 May 2. PMID: 15902306; PMCID: PMC1088017.

26. Khan FH, Shaw L, Zhang W, Salazar Gonzalez RM, Mowery S, Oehrle M, Zhao X, Jenkins T, Setchell KD, Inge TH, Kohli R. Fibroblast growth factor 21 correlates with weight loss after vertical sleeve gastrectomy in adolescents. Obesity (Silver Spring). 2016 Nov;24(11):2377–2383. doi: 10.1002/oby.21658. Epub 2016 Sep 12. PMID: 27615057; PMCID: PMC5846337.

27. Frikke-Schmidt H, Hultman K, Galaske JW, Jørgensen SB, Myers MG Jr, Seeley RJ. GDF15 acts synergistically with liraglutide but is not necessary for the weight loss induced by bariatric surgery in mice. Mol Metab. 2019 Mar;21:13–21. doi: 10.1016/j.molmet.2019.01.003. Epub 2019 Jan 14. PMID: 30685336; PMCID: PMC6407365.

28. Dolo PR, Yao L, Liu PP, Widjaja J, Meng S, Li C, Zhu X. Effect of sleeve gastrectomy on plasma growth differentiation factor-15 (GDF15) in human. Am J Surg. 2020 Sep;220(3):725–730. doi: 10.1016/j.amjsurg.2020.01.041. Epub 2020 Jan 28. PMID: 32014297.

